# Genetic regulation of injury induced heterotopic ossification in adult zebrafish

**DOI:** 10.1101/2024.02.09.579618

**Authors:** Arun-Kumar Kaliya-Perumal, Cenk Celik, Tom J. Carney, Matthew P. Harris, Philip W. Ingham

## Abstract

Heterotopic ossification is the inappropriate formation of bone in soft tissues of the body. It can manifest spontaneously in rare genetic conditions or as a response to injury, known as acquired heterotopic ossification. There are several experimental models for studying heterotopic ossification from different sources of damage. However, their tenuous mechanistic relevance to the human condition, invasive and laborious nature and/or lack of amenability to chemical and genetic screens, limit their utility. To address these limitations, we developed a simple zebrafish injury model that manifests heterotopic ossification in response to micro-fractures in combination with muscle injury. These findings indicate that clinically-emulated injuries in zebrafish can lead to osteo-induction and proliferation as observed in heterotopic ossification in *myositis ossificans traumatica*. Exploiting this model, we analysed the penetrance and expressivity of heterotopic ossification and defined the transcriptional response to trauma, identifying differentially regulated genes. Taking advantage of defined mutants in several of these candidates, we explored their impact on heterotopic bone formation. Our findings revealed that an increase in potassium channel Kcnk5b activity potentiates injury response. In contrast, we demonstrate that inflammatory responses are essential for the ectopic bone growth, as mutations in Interleukin 11 receptor paralogue (Il11ra) exhibit a drastically reduced ossification response. Based on these findings, we postulate that enhanced ionic signaling, specifically through Kcnk5b, regulates the intensity of the skeletogenic injury response, which, in part, requires immune response regulated by Il11ra.

## Introduction

The musculoskeletal system is an intricate association of muscles, tendons, ligaments, bones and joints that together support posture and enable locomotion and protection (Roberts, 2002). The state of this construction is far from static; rather, through integration of strain, localized new bone can be formed to accommodate differential strain, turnover, or repair (Bergmann et al., 2010; Watanabe-Takano et al., 2021). When these remodelling signals are aberrant or enhanced, ectopic growth of bone can result. If the enhanced bony growth involves the joints and associated structural connectivity of the skeleton, this can lead to severe effects on quality of life for patients. Research into extreme systemic heterotopic ossification such as fibrodysplasia ossificans progressiva (FOP) suggests that local signals can drive such ectopic bone formation (Pignolo et al., 2013).

FOP is a heritable disorder of the skeleton characterised by heterotopic ossification predominantly at muscles, tendons, ligaments, and fascia (Kaplan et al., 2008). Similarly, progressive osseous hyperplasia and Albright’s hereditary osteodystrophy are characterised by heterotopic ossification predominantly at cutaneous and subcutaneous sites (Kaplan and Shore, 2000). FOP has been the subject of comprehensive research and has yielded significant insights into the formation of heterotopic bone. In patients with FOP, a gain-of-function mutation in the *ACVR1* gene, encoding the Activin A receptor type 1, a bone morphogenetic protein (BMP) type 1 receptor, drives the heterotopic ossification process by initiating both ligand dependent and independent activation of the BMP signalling cascade aberrantly (Hino et al., 2015; Pang et al., 2016), a response coincident with local tissue damage and inflammation (Matsuo et al., 2019).

While these conditions are driven by underlying genetic disorders, heterotopic bone formation can also occur simply in response to triggering events, particularly injury (Meyers et al., 2019). There are two such conditions, myositis ossificans traumatica (MOT) and neurogenic heterotopic ossification (NHO), collectively referred to as acquired disorders of heterotopic ossification. MOT, being the most common type and observed primarily in young adults, typically follows a benign course and is triggered by a single traumatic incident or repeated injuries to the same anatomical site. Various musculoskeletal traumas such as fractures, muscle contusions, wartime wounds, burns, and surgeries have shown development of heterotopic ossification at the site of injury (Ahrengart, 1991; Andreu Martinez et al., 2007; Beiner and Jokl, 2002; Forsberg et al., 2009). Furthermore, repeated injuries, for example as seen in horseback riders, have been shown to lead to the development of ectopic bone, referred to as ‘rider’s bone,’ in the origin of the adductors of the thigh (Binnie, 1903; Rosenstirn, 1918). Similarly, shooters may develop ectopic bone in the deltoid muscle, known as ‘shooter’s bone (Walczak et al., 2015). Neurogenic Heterotopic Ossification (NHO) by contrast, involves the occurrence of heterotopic ossification around a fracture or a contusion site, especially around the hip joint, whenever there is a concomitant traumatic brain or spinal cord injury (Genet et al., 2009; Sakellariou et al., 2012).

These conditions involving injuries are quite instructive as the underlying mechanism of ectopic bone formation is believed to follow a particular trend. Firstly, following a muscle or bony injury there is infiltrative bleeding and inflammation (Beiner and Jokl, 2002). The subsequent cascade of events normally results in either complete anatomical healing or scar formation. However, the dysregulation of signaling of the cells at the site of injury can lead to their inappropriate differentiation into chondrocytes or osteoblasts, ultimately resulting in heterotopic bone (Kan et al., 2009). Once triggered, little can be done to alter the natural course of the disorder. Worse still, invasive procedures can enhance ongoing inflammation and lead to excessive heterotopic bone formation. Therefore, management is typically conservative and excision is only recommended after the heterotopic bone fully matures (Meyers et al., 2019). By that time, however, stiffness and muscle contractures have already developed, limiting full recovery.

Since the underlying mechanisms of both genetic and acquired heterotopic ossification are closely related, it seems plausible that common targets for intervention can be identified, allowing alleviation of growth without inducing added insult or inflammation. While current research on FOP has led to the discovery of certain drugs that are undergoing clinical trials (Pignolo et al., 2022; Pignolo et al., 2023), it remains unlikely that these would be effective for damage induced exposures, given the abnormal enhanced sensitivity seen in the gain-of-function condition of FOP etiology. In light of the higher incidence of acquired heterotopic ossification occurring in standard injuries, it is important to address the etiology of these disorders and bridge our knowledge gap concerning the mechanisms underlying the tissue response specific to this injury-induced pathology. This requires the utilization of animal models of injury induced heterotopic ossification, of which there are currently limited examples (Kan and Kessler, 2011).

Excessive manipulation of an immobilized knee in a rabbit model for a duration of 5 weeks, results in the development of heterotopic ossification along the periosteum of the femur (Bartlett et al., 2006; Michelsson et al., 1994). An Achilles tenotomy model in rats and mice, in which there is an experimental partial division of the Achilles tendon resulted in heterotopic bone formation within three months (Buck, 1953; McClure, 1983). Simulation of hip arthroplasty or femoral nailing by surgical reaming of the femoral canal in rabbits resulted in consistent heterotopic ossification at the entry site (Schneider et al., 1998). In mice, direct implantation of BMP containing matrix (BMP2 or 4) leads to the formation of abnormal bone at the implantation site (Glaser et al., 2003; Lounev et al., 2009), which may reflect the generation of non-physiological levels of osteogenic factors (Anthonissen et al., 2014). A common injury underlying heterotopic ossification involves the muscles. A simple injection of 40% ethanol or acid alcohol in rabbits, showed heterotopic bone at the injection site, primarily at muscle insertions (Heinen et al., 1949; Selle and Urist, 1961). In dogs, stripping the periosteum and damaging the overlying muscles of the femur resulted in heterotopic ossification, further enhanced by repeated blunt trauma to the injury site (Collins et al., 1965). Similarly, sheep showed heterotopic ossification within 3 weeks to 3 months following a crush injury to the muscles overlying the femur (Walton and Rothwell, 1983). While some of these models replicate specific modes of heterotopic ossification occurrence in humans, others lack mechanistic relevance, or their procedures are invasive and laborious, making them unsuitable for broad-scale experimental analysis. An established and broadly applicable injury-induced heterotopic ossification model is thus still lacking.

Zebrafish are amenable to experimental manipulation, such as surgical extirpation as well as damage models in the study of repair and regeneration, and as such present an attractive model for the development of an experimental platform for heterotopic ossification. Notably, the powerful genetics in zebrafish facilitates investigation of the genetic and cellular activity underlying these responses. There are remarkable parallels between zebrafish and mammals in their musculoskeletal development and remodelling. In particular, zebrafish skeletal cells, the patterns of ossification, and the sequential transcriptional hierarchy driving osteogenesis share striking similarities with those observed in mammals (Kaliya-Perumal and Ingham, 2022). In the last decade, the zebrafish has become an important organism for the study of skeletogenesis, particularly in mature tissues, serving as models of skeletal dysplasia such as osteogenesis imperfecta, osteopetrosis, and a broad range of structural birth disorders (Mari-Beffa et al., 2021). The zebrafish has also been used to model the genetic signalling in FOP (Allen et al., 2020; LaBonty et al., 2017; LaBonty et al., 2018).

Here we investigate the efficiency and character of experimental induction of heterotopic ossification in the adult zebrafish, specifically to emulate mechanistically relevant injuries causing these disorders in humans. We capitalize on this model to assess genetic regulation of this response and novel targets for its intervention.

## Results

### Exploring heterotopic ossification

Zebrafish has become an established model system for the analysis of human disease genetics and within the last decade has been increasingly exploited as a model for disease that affect adult structures, including the skeleton (Carnovali et al., 2019; Valenti et al., 2020). We postulated that by employing relevant injury methods, heterotopic ossification could manifest in zebrafish adults. To pursue this, we chose the caudal peduncle region just rostral to the tail fin as a site wherein mechanistically relevant contusion or fracture-contusion injuries could be inflicted, mirroring those occurring in humans. This region of the fish is muscular, remote from essential organ functions, and adjacent to articulation of the tail fin bones with the hypurals and parahypurals. In addition, this area favours live fluorescence imaging of the skeleton utilising transgenic reporters such as *Tg(sp7:gfp*), which drives GFP expression in the early osteoblasts throughout the body. Hence, the entire skeleton exhibits bright green fluorescence including heterotopic ossification sites starting from the early stages.

A lancing device was designed for the controlled induction of initial contusions. Observing resolution of singular damage events within 48 hours in all the fish, the injury site was gently stroked with a blunt-tip Kirschner pin (K-pin) to revert it to its contused state. This process was repeated, sustaining the contusion phase for up to a week. After a 4-week interval, live imaging was carried out (Fig 1). The tail fin lepidotrichia were clearly visible, extending to the proximal region beneath the overlying soft tissue of the caudal peduncle (Fig 1B). However, the hypural, para-hypural, and the connection points of these bones with the fin rays remained obscured by the surrounding soft tissue cover and were not visualized. Upon staining and imaging, new bone formation resembling heterotopic ossification was observed at the caudal margin of the contusion site, originating from the lepidotrichia beneath the contused muscle and occupying the adjacent soft tissue area. While this site falls outside the trajectory of the lancet strikes, the resulting contusions spread across a wider area, including the region where ectopic bone is noted. In one of these fish, robust bridging bone formation was noted, connecting multiple adjacent fin rays (Fig. 1E). However, the overall occurrence rate was found to be minimal (26.1%, n=23) (Fig 1D). Hence, it became imperative to identify a more suitable and pertinent location with higher penetrance for further research.

**Fig. 1.**
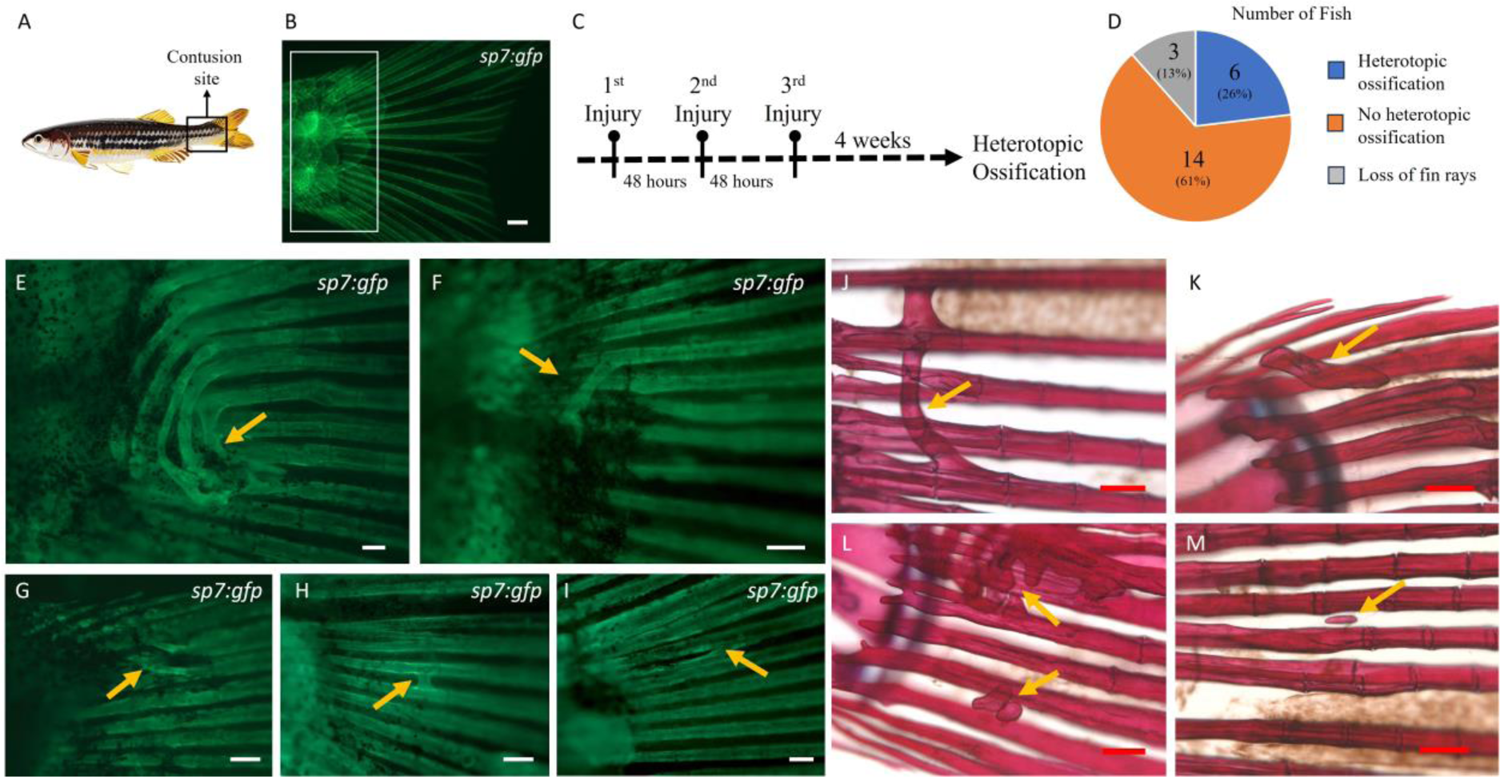
Heterotopic bone formation at the tail fin lepidotrichia. A) Site of caudal peduncle contusion (boxed area). B) Appearance of an uninjured tail fin under fluorescent microscopy in a transgenic *Tg(sp7:gfp)* zebrafish. The boxed area represents the site where heterotopic bone formation was noted, as shown in E-M. C) Timeline representing the recurring injuries with a 48-hour interval in between, followed by a 4-week wait to observe heterotopic bone. D) Pie chart showing the overall number of fish with and without heterotopic bone (n = 23). E) Exuberant heterotopic bone formation (arrow) bridging between multiple adjacent fin rays beneath the overlying soft tissues of the caudal peduncle. F–M) Various other forms of heterotopic bone (arrows) observed. Scale bars: B, E – 500µm; F – M – 200µm.

### Intermuscular bone hypertrophy at contusion sites

Intermuscular bones form an integral part of the skeletal system in teleost fish. They are located in the myosepta on both sides of the vertebrae and develop via intramembranous ossification of myoseptal tendons without undergoing a cartilaginous phase (Fig 2C, D) (Nie et al., 2019). In most teleost fish, there are three sets of intermuscular bones, epineurals, epicentrals and epipleurals attaching ligamentously to neural, central and hemal arches or ribs, respectively (Nie et al., 2019). In the zebrafish, however, only epineural and epipleural bones are present, the former in each myoseptum and the latter only in myosepta caudal to the ribs throughout the tail region (Bird and Mabee, 2003; Yao et al., 2015). Their function is to transmit force between muscle segments and increasing the stiffness of the body. During development of the zebrafish, these bones ossify from distal to proximal, and the ossification is believed to be influenced by the mechanical load induced by swimming (Yao et al., 2015).

Following caudal peduncle contusion, we observed a larger-than-normal size in the hypurals, parahypural, as well as the corresponding haemal and neural spines (Fig 2E). The increase in bone size was evident when comparing them to the typical appearance of these bones in control fish of the same age and size. Furthermore, we observed that the intermuscular bones located at the injury site, specifically the distal most and pre-distal pairs, which are entirely encased within muscle tissue and not connected to the axial spine, displayed signs of proliferation and enlargement in the majority of the injured fish (Fig 2F). This observation raises the possibility that the muscle contusion alone could trigger factors conducive to osteoproliferation.

To examine a similar occurrence at an alternate, non-mobile site, we induced recurring unilateral contusions by gentle stroking using a blunt tip K-pin on the dorsal thoracic region, designating the uninjured side as the control. We observed a striking enlargement of the intermuscular bones on the contused side in all the injured fish (Fig 2H, I). In addition, some of the fish demonstrated hypertrophy of the neural spines in the region (Fig 2J). This observation confirms the osteo-proliferative characteristic of the contusion site. To quantify the changes, we outlined the bones and measured the contained area. This process was repeated for the corresponding uninjured side, and the extent of hypertrophy was determined by comparing the bones on either side. It was evident that the intermuscular bones (n = 12) were significantly larger on the injured side (Uninjured = 13,807.01 ± 3385.35 µm^2^; Injured = 57,731.8 ± 18801.4 µm^2^) by more than four-fold (4.18; p<0.0001) (Fig 2K). Given that osteo-induction and subsequent proliferation is required for development of any bone, the zebrafish injury microenvironment harbouring these properties following injury can serve as a valuable tool to study heterotopic bone formation.

**Fig. 2.**
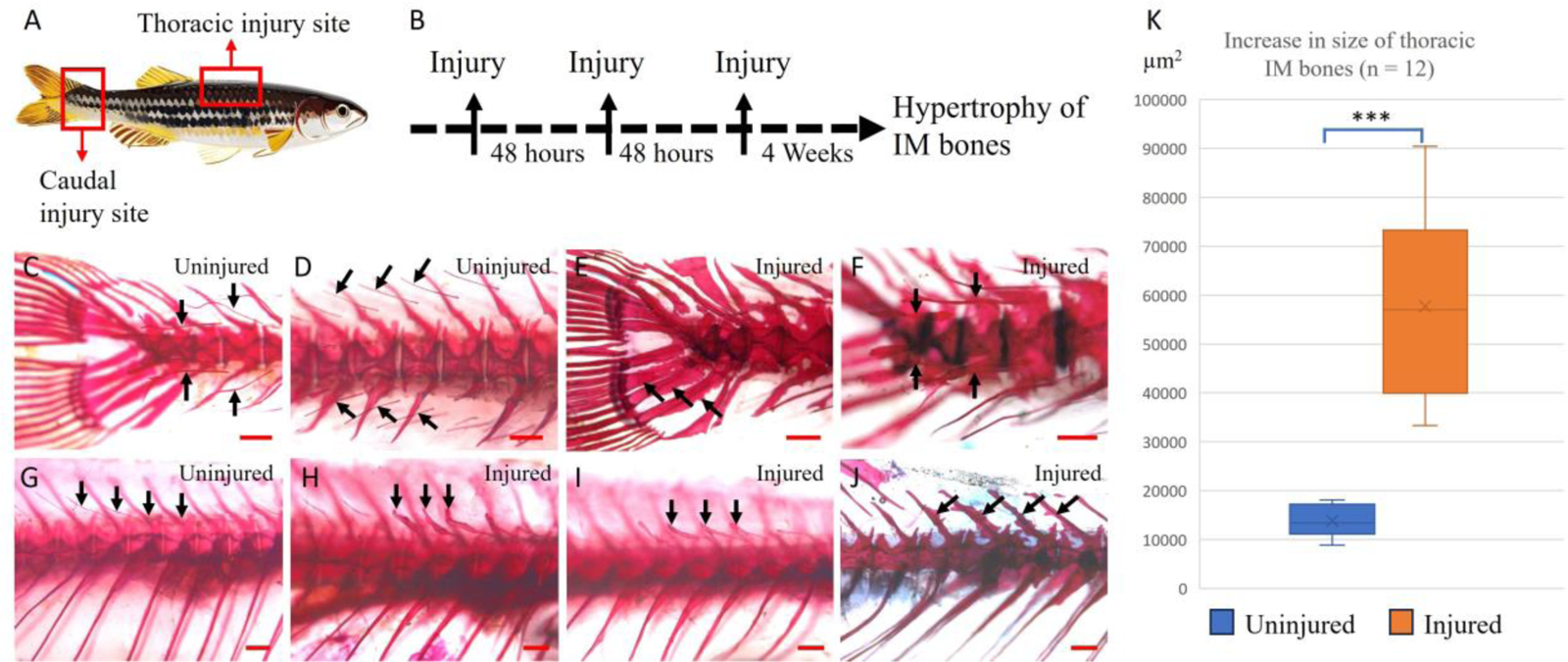
Hypertrophy of intermuscular bones following contusions. A) Boxed areas represent the contusion sites. B) Injury timeline, similar to pectoral fin injuries but resulting in hypertrophy of intermuscular (IM) bones. C, D) Uninjured caudal peduncle sites showing the intermuscular bones (arrows). E) Injured caudal peduncle site, one month later, showing hypertrophied hypurals and parahypural (arrows). F) Injury site showing hypertrophied intermuscular bones (arrows). G) Uninjured thoracic region showing normal epineural intermuscular bones (arrows). H, I) Injured thoracic region, one month later, showing hypertrophied intermuscular bones (arrows). J) Hypertrophied neural spines in an injured fish (arrows). K) Graph showing the significant size increase observed in the intermuscular bones on the side of injury compared to the uninjured side (Y-axis is in μm²). Scale bars: 500 µm.

### Identification of highly penetrant model for heterotopic bone involving the pectoral fin

The medial aspect of the pectoral fin was selected as an alternative site due to its mechanistic relevance. In this region, the bulk of muscles responsible for fin movements, notably the *adductor superficialis* and *adductor profundus*, envelop the scapulo-radialis bones and the proximal section of the fin rays (Siomava and Diogo, 2018). In addition, the fin rays here are bulkier and stronger than those at the tail fin. Here, we induced heterotopic bone by creating injuries involving both muscle and bone near the articulation of the pectoral fin with the radial bones, precisely where the dorsal hemi-ray and ventral hemi-ray fuse (Fig 3B).

Fish were anaesthetised and the pectoral fins were injured as described in Methods. Injury to the medial muscle bulk and microscopic damage to the proximal aspect of the pectoral fin rays were observed post-injury (Fig 3E, F). Similar to the caudal peduncle injuries, injuries were repeated twice at 48-hour intervals to sustain inflammation. Live imaging of this region posed challenges because the body of the fish beneath the pectoral fin obstructs the optical plane. However, using double transgenic *Tg(sp7:gfp;ctsk:dsred)* fish, we identified early macrophage and osteoclastic activity in response to the damage (Fig 3C, D, G, H). The fish were stained at one month for Alizarin red staining and analysis of heterotopic bone. All fish exhibited heterotopic bone formation, manifesting as spurs and/or bridges between the rays (n=5, Fig 3I-M). This penetrance was consistently reproducible in subsequent batches of fish with distinct heterotopic bone formation observed in all the injured fish (n=32). As generating these injuries is straightforward and the formation of the pathology is fully penetrant, this model is easily scalable and reproducible for studying heterotopic ossification.

**Fig. 3.**
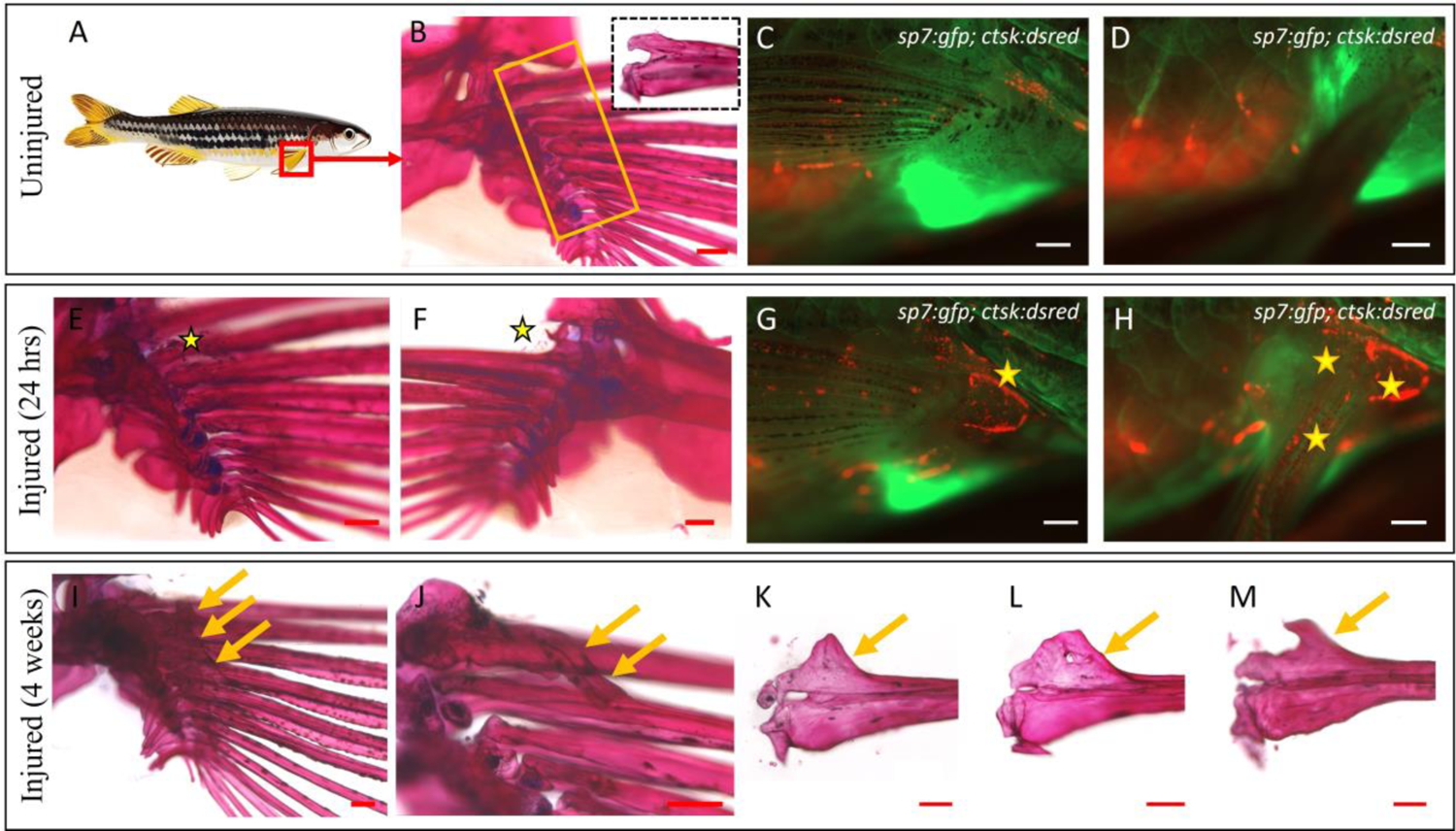
Consistent heterotopic ossification in the form of spurs and bridges in the pectoral fin. A) Graphic depicting the injury site. B) Medial view of an uninjured right pectoral fin stained with alizarin red; yellow box indicates the injury site; zoomed in dotted box focuses on the articular region of a dissected fin ray, comprising the dorsal and ventral hemirays. C) Lateral view of an uninjured pectoral fin of *Tg(sp7:gfp; ctsk:dsred)* fish under fluorescence. D) The same fish after elevating the pectoral fin to show the inner (medial) side where the absence of osteoclastic activity can be noted. E, F) Medial and lateral views of an injured pectoral fin, 24 hours post injury, showing tiny flecks of bone (star) arising from the microfractures. G) Lateral aspect of an injured pectoral fin of *Tg(sp7:gfp; ctsk:dsred)* fish showing osteoclastic activity (star) at 24 hours post injury. H) Osteoclastic activity on the medial aspect seen after elevating the fin (stars). I) Injured pectoral fin, one month after injury. Arrows highlight the heterotopic bone spurs. J) Heterotopic bridging bone noticed between the marginal ray and the 2nd ray in one of the injured fish (arrow). K-M) Different forms of heterotopic bone spurs encountered following injury (arrows). Scale bars: B, E, F, I - M – 200µm; C, D, G, H - 500µm.

### Bulk RNA Sequencing for transcriptional profiling of injured tissue

To investigate the transcriptional response underlying the observed proliferation and enlargement of bones at the contusion site, we conducted genome-wide bulk RNA sequencing on contused muscle from the caudal peduncle region, as this region allows for easy harvesting of the entire muscle bulk. Investigation was not solely focused on comparing uninjured and injured conditions, but rather assessed the response characteristics in the case of multiple recurring injuries. Contused muscle tissue was dissected at four different time points: No injury (Control), 24 hours after single injury (SI), 24 hours after multiple (three) injuries (MI) and 5 days after multiple (three) injuries (D5MI). Principal component analysis (PCA) analysis revealed similarity between replicates and differences across conditions (Fig 4B). Further analysis was focused on three comparisons: 1) SI vs Control, 2) MI vs Control and 3) D5MI vs Control where Benjamini-Hochberg adjusted *p* values below 0.05 and log2FC of 1.5 were considered significant for determining differentially expressed genes.

The numbers of upregulated and downregulated genes depicted a combination of both unique and overlapping signatures per condition (Fig 4C, D). Pairwise comparisons revealed 1193 up and 1849 down regulated genes for the SI, 2192 up and 2229 down regulated genes for the MI and 2286 up and 1873 down regulated genes for the D5MI conditions (Fig 4E-G). Assessing the top 10000 differentially expressed genes (DEGs) to understand the global transcriptomic changes revealed a temporal relationship with expression profiles of the D5MI group more closely related to the MI group when compared to the SI group and control (Figure S1). Analysis of the DEGs (Table S1-S3), Gene Ontology (GO) classification and pathway enrichment analysis (KEGG), suggested that immune response characterised by apoptosis and upregulation of inflammatory mediators is triggered after muscle contusion injury (Figure S2-S4). As a result of recurring injuries, this immune signature persisted. Specifically, subsequent to an initial upregulation of genes *il11a* and *il11b,* an elevation in various osteoblast differentiation markers was observed over time. These include *runx3, bglap, dlx* transcription factors*, msx2a, spp1, fn1a,* along with teleost-specific genes *scpp1, scpp5,* and *scpp7* (Huang et al., 2007; Kawasaki, 2009). In addition, developmental patterning genes such *sall1a* and *sall1b*, *hoxa13a* and *hoxa13b,* (Perez et al., 2010) were upregulated. Interestingly, also upregulated were *kcnk5b,* which acts locally within the mesenchyme of fins and barbels to specify appendage size (Perathoner et al., 2014), and osteoclast specific *stat1b, ocstamp, csf1ra* and *ctsk*, though at the later time point (D5MI) (Fig 4I). These notable changes could underlie the proliferation and increase in size of bones after injury.

**Fig. 4.**
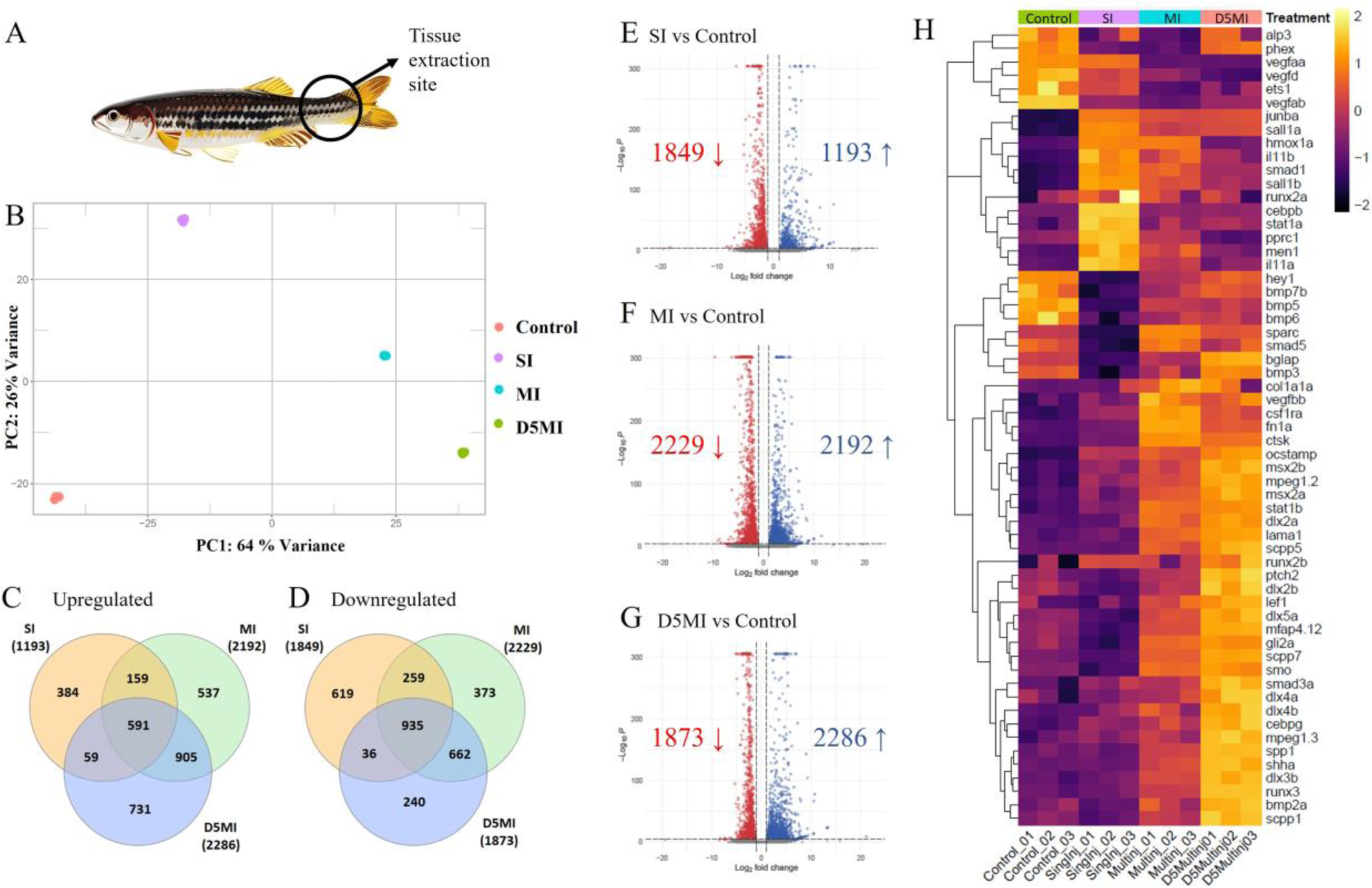
RNAseq cluster analysis and differential gene expression. A) Graphic showing the caudal peduncle contusion site from where tissues were extracted. B) Principal component analysis (PCA) plot displaying sample distances. C, D) Venn diagrams illustrating the number of upregulated and downregulated genes in each comparison. E-G) Volcano plots depicting the overall number of upregulated and downregulated genes among the three comparisons, with log fold change on the X-axis and significance on the Y-axis. H) Heat map of selected genes of interest (SI: 24 hrs after single injury; MI: 24 hrs after multiple injuries; D5MI: 5 days after multiple injuries).

### The role of potassium channel activity in regulating response characteristics of heterotopic bone formation

It has been shown that elevated activity of potassium channels can influence skeletal growth and patterning in zebrafish (Perathoner et al., 2014). These channels are crucial for regulating cellular electrical potential and are involved in diverse biological processes, but their precise activity in tissues is not well characterized or understood. Examining *kcnk5b* expression in response to contusion injuries in our transcriptome data, we observed a progressive and significant increase over-time (Fig 5A). This observation was further validated through qRT-PCR analysis (Fig 5B). Consequently, we were interested in exploring the heterotopic bone formation response in the context of altered Kcnk5b activity. The *pfau* gain-of-function mutants of Kcnk5b display elongated fin segments due to increased growth rate of lepidotrichia (Perathoner et al., 2014). We observed an accelerated response in injured fish (n = 4), with detectable heterotopic bone formation occurring in just two weeks after the last injury, in contrast to response of wild-type fish (Fig 5D, E; Movie 1). In a separate batch, one month after the injury, all affected pectoral fins (n = 5) exhibited markedly greater heterotopic bone formation compared to their wild-type counterparts (Fig 5G, H). Notably, in one of the injured pectoral fins, we observed bridging between the fin rays and the radial bones (Fig 5H), emphasizing the extensive nature of the heterotopic response. To quantify these findings, we stained the fins with 1% silver nitrate and performed micro-CT scans (Fig 5I-L) (Charles et al., 2017). Using 3D Slicer image computing platform, we compared the bone volume of the injured and uninjured side in all the fish. Our findings reveal that the *kcnk5b^pfau^*^+/-^ mutants, in contrast to wild-type fish, exhibit a significant (p = 0.02) increase in bone volume at the injured pectoral fin by two weeks following the last injury. Furthermore, after one month, while the wild-type fish showed a 17.1±1.5% increase in bone volume on the injured fin, the *kcnk5b^pfau^*^+/-^ mutants displayed a significantly greater increase of 33.6±11.9% (p = 0.01) (Fig 5M). Therefore, it appears likely that potassium channel signalling, specifically via Kcnk5b, plays a crucial role in both initiation and manifestation of the heterotopic bone formation response.

**Fig. 5.**
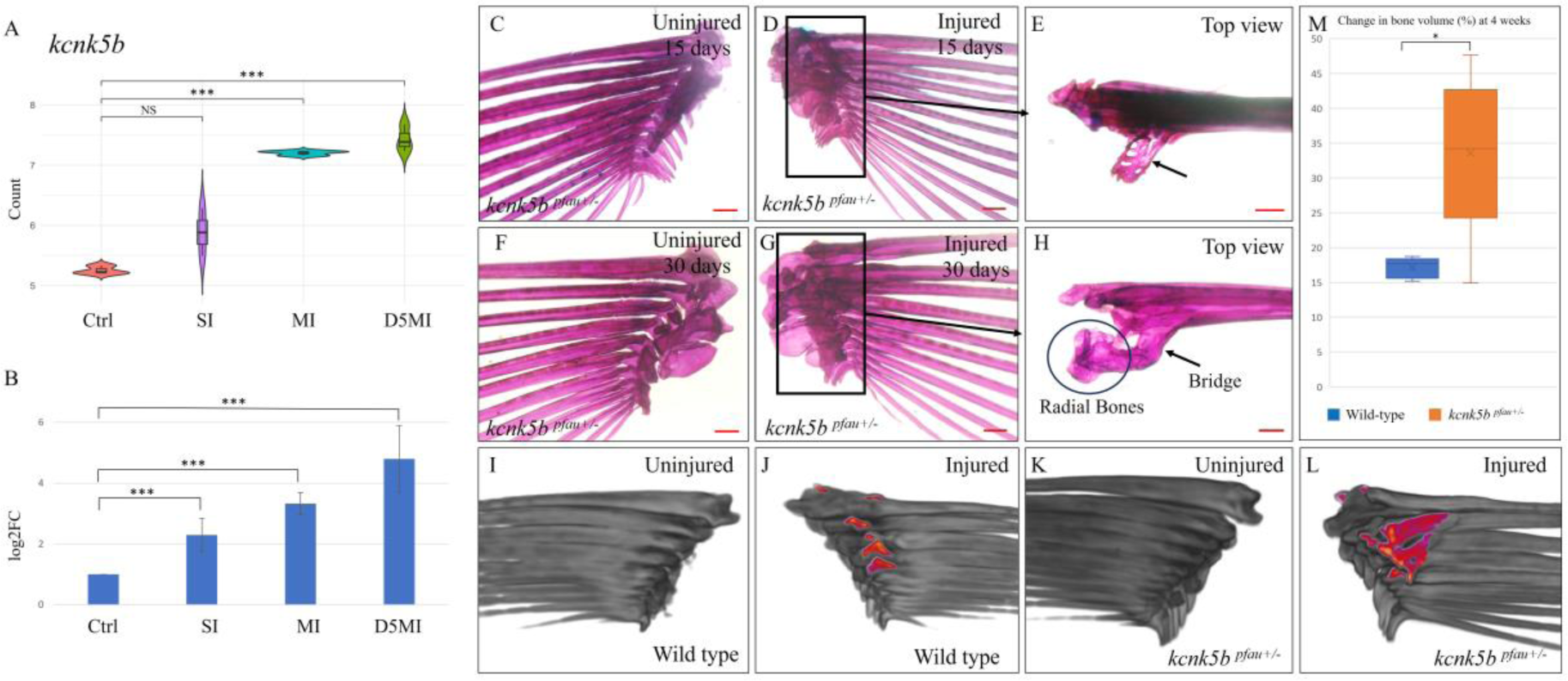
Differential expression of *kcnk5b* and higher magnitude of heterotopic bone formation in *kcnk5b^pfau^*^+/-^ mutants. A) Violin plot depicting the differential expression of *kcnk5b* as inferred from RNA sequencing. B) qRT-PCR validation of the same. C, D) Uninjured left and injured right pectoral fins of a *kcnk5b^pfau+/-^* mutant zebrafish visualized 2 weeks post-injury. D) Medial aspect of the injured pectoral fin showing extensive heterotopic ossification (boxed area). E) Top view of the same clearly illustrating the extent of heterotopic bone. F, G) Uninjured left and injured right pectoral fins of a *kcnk5b^pfau+/-^* mutant zebrafish visualized one-month post-injury. G) Medial aspect of the injured pectoral fin showing extensive heterotopic ossification (boxed area). H) Top view of the same indicating bridging (arrow) between the 2nd fin ray and the radial bones. I-L) 3D reconstructed CT scans. I, J) Medial view of the uninjured left and injured right pectoral fins of a wild-type fish. J) Injured pectoral fin showing heterotopic bone spurs highlighted in colour. K, L) Medial view of the uninjured left and injured right pectoral fins of a *kcnk5b^pfau+/-^*mutant fish. L) Injured pectoral fin showing extensive heterotopic bone (highlighted in colour) unlike wild-type fish. M) Box and whisker plot showing a significant increase in bone volume following injury, unlike in wild-type fish. *p < 0.05, ***p < 0.001. Scale bars: C-H - 200µm.

### Role of IL-11 signalling in the development of heterotopic ossification

Expression of *il11a* and *il11b* was significantly upregulated as an early response to contusion injuries, the levels of both transcripts gradually declining at later time points (Fig 6A, B). This observation was validated through qRT-PCR analysis (Fig 6C, D). Previous studies revealed that Il-11 encoding genes (*il11a* and *il11b*) exhibit the most significant induction and highest expression after tissue damage in the zebrafish cardiac ventricle and caudal fins (Allanki et al., 2021). Moreover, transcriptome data from various regenerating tissues in zebrafish, African killifish, lungfish, *Xenopus*, and axolotl have demonstrated an evolutionarily conserved and injury-responsive induction of Il-11 (Darnet et al., 2019; Fang et al., 2013; Gerber et al., 2018; Tsujioka et al., 2017; Wang et al., 2020). In the case of zebrafish, mutants lacking the Il-11 receptor, *il11ra* gene, similar to *Il11r* mutant mice, survive into adulthood without noticeable developmental defects (Allanki et al., 2021). Upon injuring these mutants (n = 5), it was observed that the *il11ra*^-/-^ mutants failed to exhibit heterotopic bone formation similar to their wild-type counterparts (Fig 6G, I; Movie 2). Notably, while wild-type siblings exhibited a significant increase in bone volume in injured pectoral fins, the *il11ra*^-/-^ mutants, in contrast, demonstrated a significant reduction (-12.4±5.9%; p<0.0001) in bone volume (Fig 6J). This difference was evident both visually and in volumetric assessments using micro-CT scans. Although we noticed a single spur formation in one of the injured *il11ra*^-/-^ mutants, the overall bone volume in the injured fin was lower compared to the uninjured side. These observations emphasize the pivotal role that IL-11 signalling plays in the normal damage response program. Furthermore, they suggest that targeting of IL-11 function and its suppression may have implications for reducing injury-induced inflammation and the subsequent development of heterotopic bone.

**Fig. 6.**
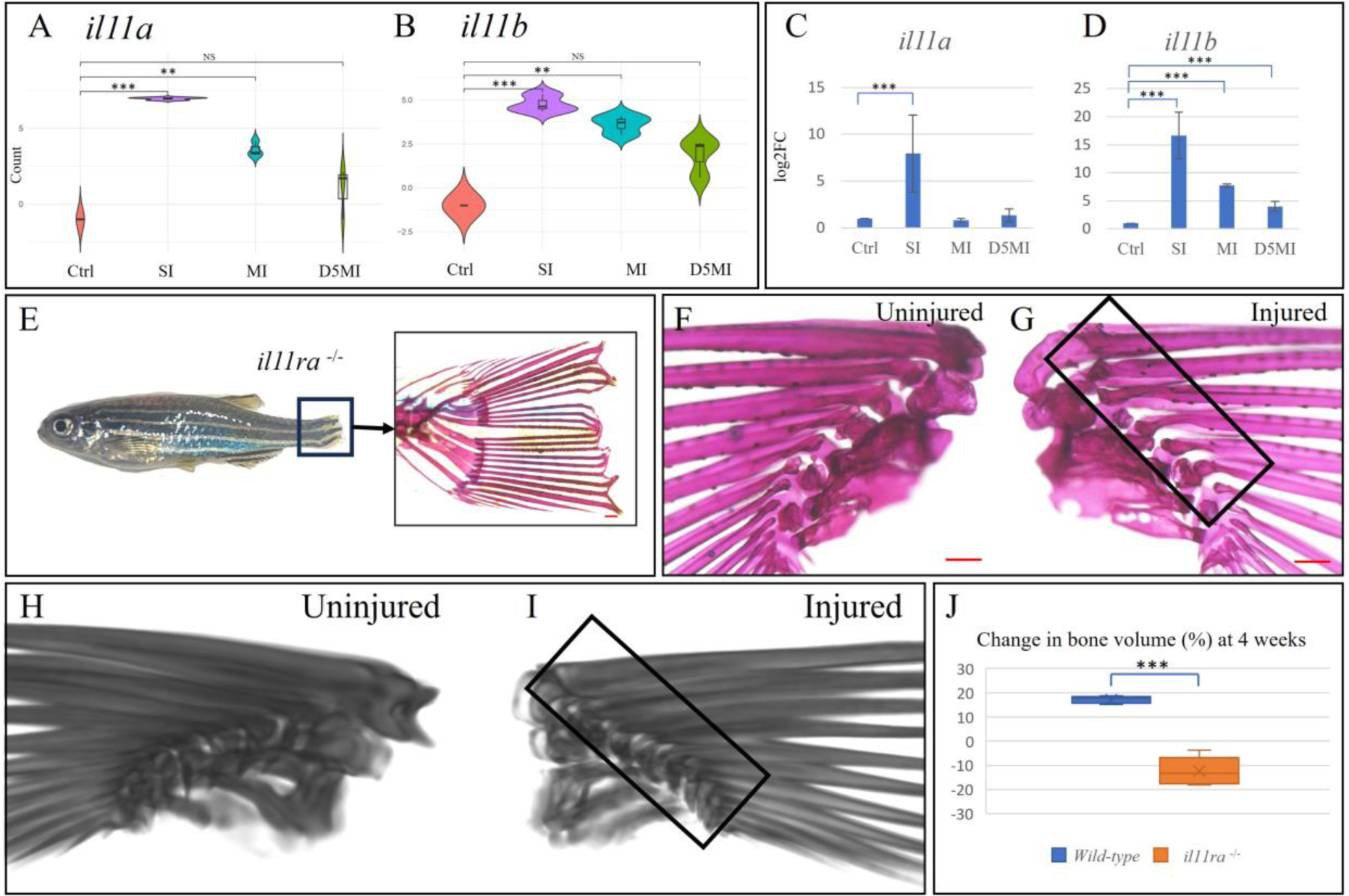
Differential expression of genes encoding Il-11, and absent heterotopic ossification in *il11ra^-/-^* mutants. A, B) Violin plots depicting the differential expression of *il11a* and *il11b* as per RNA sequencing. C, D) qRT-PCR validation of the same. E) *il11ra*^-/-^ mutant zebrafish demonstrating impaired tail fin regeneration (boxed area) at 2 weeks following injury. Zoomed-in image shows an alizarin red-stained section of a non-regenerated tail fin. F, G) Uninjured left and injured right pectoral fins of an *il11ra*^-/-^ mutant zebrafish visualized one-month post-injury. G) Medial view of the injured right pectoral fin showing no signs of heterotopic bone. H, I) 3D reconstructed CT scans. I) Medial view of the injured right pectoral fin shows no heterotopic bone. I) Box and whisker plot showing a significant reduction in bone volume following injury, unlike in wild-type fish. **p < 0.01, ***p < 0.001. Scale bars: E-G - 200µm.

### Intermuscular bone hypertrophy in *kcnk5b* gain-of-function and *il11ra* loss-of-function mutants

Given the heterotopic bone formation responses observed in fish carrying either the *kcnk5b* gain-of-function mutation or *il11ra* loss-of-function mutation, we explored the development of intermuscular bone hypertrophy resulting from thoracic contusions in both mutants. We assessed intermuscular bone hypertrophy response conducted in a series of three injury episodes, each separated by a 48-hour interval as demonstrated previously, using Alizarin red staining. Our findings revealed an increase in the size of intermuscular bones at the injury site in all fish, regardless of their mutation status (Fig 7A). However, when we examined these changes more closely, we observed distinct differences between the mutant and wild-type fish. Specifically, fish carrying the *kcnk5b* gain-of-function mutation exhibited a notably robust increase in intermuscular bone size, highlighting a significant response to the injury (Fig 7B). On the other hand, fish homozygous for the *il11ra* loss-of-function mutation displayed a more limited increase in intermuscular bone size. These findings provide valuable insights into the role of these mutations in the response to thoracic contusions and subsequent intermuscular bone hypertrophy.

**Fig. 7.**
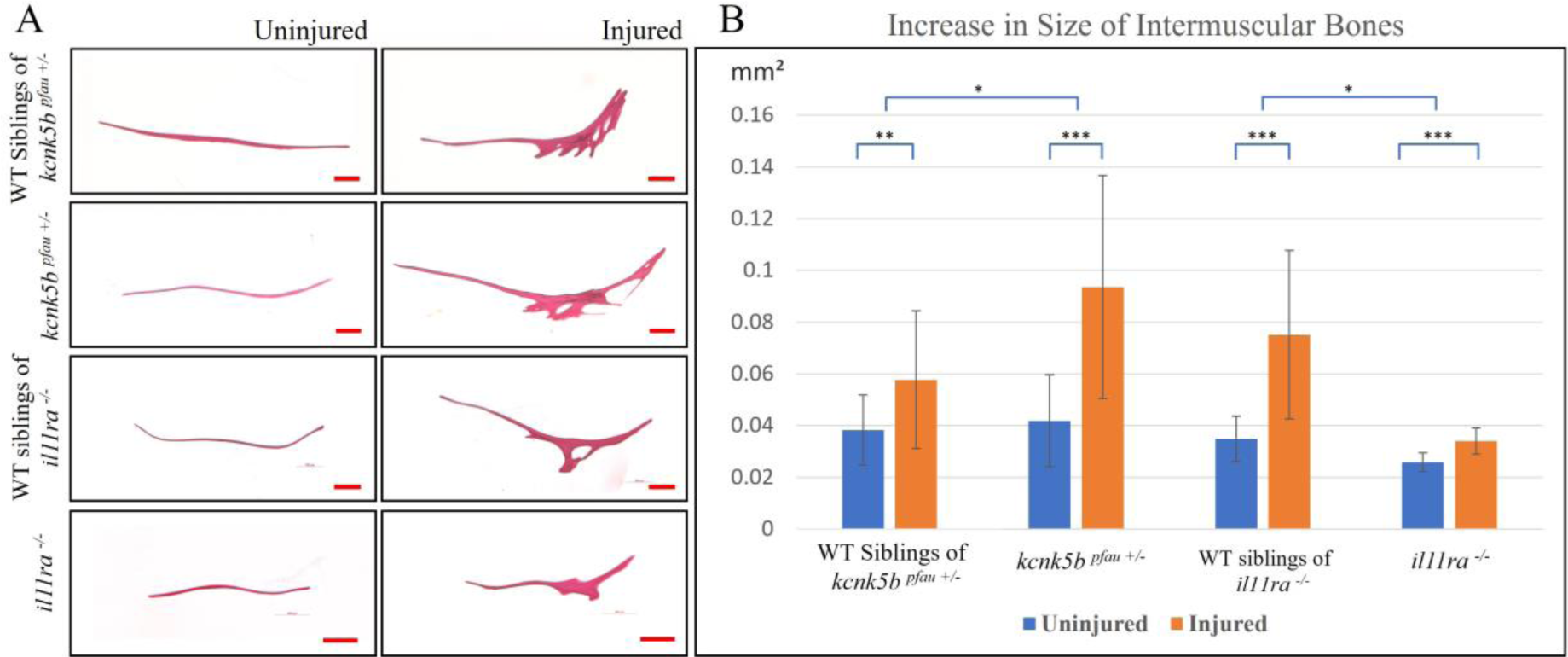
Analysis of intermuscular bone characteristics following injury in *kcnk5b^pfau+/-^* and *il11ra^-/-^* mutants. A) Representative images comparing the thoracic intermuscular bones on the injured and uninjured sides in *kcnk5b^pfau+/-^, il11ra^-/-^*, and wild-type sibling controls. B) Quantitative analysis reveals a significant increase in the size of intermuscular bones in all injured fish. Additionally, a greater increase in *kcnk5b^pfau+/-^* and a limited increase in *il11ra*^-/-^ can be observed. *p < 0.05, **p < 0.01, ***p < 0.001. Scale bars: 200µm.

## Discussion

We describe the establishment of the first zebrafish model of heterotopic ossification that closely resembles human Myositis Ossificans Traumatica (MOT). The injuries carried out in this study were designed to be akin to the types of injuries occurring in humans, such as contusions and microtrauma to bone that have the potential to lead to heterotopic ossification. In contrast to previous models in mammals, these injuries in zebrafish do not necessitate a sophisticated experimental setup. Furthermore, the formation of mature heterotopic bone is rapid, occurring within a range of 3-4 weeks. The presentation of heterotopic bone also mimics numerous instances of MOT reported in the literature, where the heterotopic bone frequently originates from the underlying bone and occupies the adjacent soft tissue area. Additionally, the bridging heterotopic ossification observed between two underlying bones in some of the injured fish mimics extensive cases of heterotopic bone formation observed in patients with FOP and in some reported cases of MOT (Kanthimathi et al., 2014; Kotb et al., 2023).

There is a well-entrenched grading system for heterotopic ossification occurring after hip replacement/arthroplasty surgeries, known as the Brooker classification of heterotopic ossification (Brooker et al., 1973). According to this system, there are four grades of heterotopic ossification that can occur following hip replacement: Grade 1: Islands of bone in the soft tissues; Grade 2: Bone spurs, > 1 cm gap between opposing bony surfaces; Grade 3: Bone spurs, < 1 cm gap between the opposing surfaces; Grade 4: Complete ankylosis (Brooker et al., 1973). Examining the appearance of heterotopic bone encountered in zebrafish, we correlate responses with the aforementioned classification system, where the formation of spurs aligns with Grades II and III, and complete bridging aligns with the most severe Grade IV.

Although the skeletal structures in the zebrafish are neither homologous, nor of orthologous developmental origin (endoskeletal vs intramembranous) to human bone, our zebrafish model of pectoral fin heterotopic ossification can nevertheless be related to hip arthroplasty-induced heterotopic ossification. During hip arthroplasty, raw surfaces on the bones are generated both at the femoral end and the acetabular end. Coincidently, this is where heterotopic bone spurs often form. In the zebrafish pectoral fin, post-injury, microscopic damage to the bone was evidenced, leaving behind small fragments of bone in the soft tissue, reminiscent of the reaming debris in human surgery. While our analysis suggest this debris is resorbed by increased osteoclast activity, it results in a raw area on the surface of the bone where heterotopic ossification develops. This phenomenon can also be related to femoral intramedullary nailing, a common surgical procedure to stabilize femoral shaft fractures in humans (White et al., 2011). Generally, there are no issues; however, in rare circumstances, heterotopic ossification may be observed at the entry site, typically manifesting as large bony outgrowths attached to the underlying bone (Botolin et al., 2013; Marks et al., 1988). These reports, consistent with our model, indicate that raw surfaces on bone, even in the absence of a complete fracture, have the potential to form heterotopic bone under appropriate conditions.

The occurrence of heterotopic ossification varies in its severity and location across different types of trauma (Meyers et al., 2019). It manifests in approximately 30% of patients following fractures or dislocations in the elbow (Hong et al., 2015). High energy extremity trauma, TBI or SCI, and other neurological disorders, are reported to increase this incidence to over 50% (Forsberg et al., 2009). The highest reported incidence is associated with severe traumatic amputations, exceeding 90% (Daniels et al., 2018), and the lowest reported incidence is linked to burn injuries, ranging from 3.5% to 5.6% (Hu et al., 2021). In our model, the incidence of heterotopic ossification due to contusion injury at the caudal peduncle region was 26%. Even though this aligns with the incidence of heterotopic ossification in humans, it does not provide the needed penetrance for an efficient experimental model as is also the case with some of the previously proposed animal models of trauma-induced heterotopic ossification (Anthonissen et al., 2014; Walton and Rothwell, 1983). Alternatively, the pectoral fin following injuries demonstrated 100% penetrance, as did the intermuscular bone hypertrophy resulting from muscle contusions. Consequently, these models are suitable for scaling up to facilitate extensive experimental analysis of management strategies for heterotopic ossification disorder.

Data from patients with FOP and *in vivo* animal models of FOP suggest that the inflammatory microenvironment harbours mesenchymal stem cells (MSCs) that ultimately differentiate into osteoblasts leading to new bone formation (Billings et al., 2008; Kaplan et al., 2011). Several studies have sought to determine the source of these MSCs. Initially, cells displaying disrupted BMP signalling and abnormal osteogenic differentiation were believed to be from the myogenic lineage (Katagiri et al., 2018), however, further investigations suggested the local MSC population at the site of inflammation to be a more relevant source of progenitor cells that differentiate into chondrocytes and osteoblasts (Billings et al., 2008). Furthermore, various sources such as the local stromal/fibroblastic cells, endothelial cells through endothelial-mesenchymal transition, Scx+ tendon progenitor cells, bone marrow-derived muscle-resident Mx1+ cells, glutamate transporter (Glast or SLC1A3)-expressing progenitor cells, and certain circulating osteogenic precursor cells with access to bone-forming sites are linked to the origin of these MSCs (Dey et al., 2016; Pignolo and Kassem, 2011; Pulik et al., 2020; Ranganathan et al., 2015; Wosczyna et al., 2012). Nevertheless, this continues to be an area under continuous investigation. Considering these mechanisms, inhibiting osteoblastic differentiation may appear to be a feasible therapeutic approach. However, it cannot be executed in situations where there is a fracture, as doing so would affect fracture healing. Hence, there is a need for alternative targets.

Our data on the transcriptional response to trauma revealed gene expression signatures which have not been previously explored within the context of heterotopic ossification. There is a scarcity of published transcriptome datasets for contused muscle tissue, particularly those examining repeated injuries at multiple time points as in this study, although Ren et al. conducted a study wherein one-time muscle injuries were induced by dropping a 500 g weight from a height of 50 cm onto the right hind limb of rats using a free fall motion (Ren et al., 2021). Rats from the experimental groups were sacrificed at intervals of 4, 8, 12, 16, 20, 24, or 48 hours after the injury, and RNA sequencing was carried out on the extracted injured tissues. Our observations are in accordance with these findings, specifically when examining the outlined GO terminologies. Notably, transcripts associated with biological processes such as response to stimuli, oxidative stress, inflammation, infiltrating immune cells, apoptosis, hemopoiesis, and microtubule processes show similarity, especially during the early timepoints. In addition, our dataset displayed comparable enrichment in KEGG pathways associated with the immune response, cascade reactions, apoptosis and phagocytosis, as well as repair-related processes.

Given that the caudal peduncle contusion site exhibited heterotopic ossification at a limited frequency, it showed a more pronounced hypertrophy of intermuscular bones following injury. Notwithstanding the varied expressivity, the RNA sequencing findings closely correspond to the changes causing proliferation and increase in size of intermuscular bones. This sheds light on the differentially regulated genes at contusion sites with the potential to promote osteogenesis, despite the absence of bony injuries. Although the master switch governing this process remains to be defined, these genes and pathways can be designated as targets to be studied using genetic manipulations and pharmacological interventions to determine whether they exert any inhibitory effect on the heterotopic bone formation response.

We first focused our analysis on the observed differential expression of Kcnk5b, a two-pore potassium-leak channel that regulates membrane potential by outward flow of potassium from the cell (Goldstein et al., 2001). Although the exact mechanisms through which these ion channels modulate various pathways are not yet fully understood, their relationship is apparent. Mutations in the Kir2.1 potassium channel have been linked to Andersen-Tawil Syndrome (Ozekin et al., 2020; Plaster et al., 2001; Yoon et al., 2006). Kir2.1 plays a crucial role in bone morphogenetic protein (BMP) signalling, and genetic disruptions in this channel result in reduced activation of downstream BMP targets (Belus et al., 2018). This is linked to the modulation of BMP release via the regulation of membrane potential and the levels of intracellular calcium. In mice, the genetic removal of Kir2.1 mirrors the phenotypes seen in BMP2/4 mutants. This includes the development of severe craniofacial features such as an enlarged fontanelle, underdeveloped mandible, nasal bone hypoplasia, and cleft palate, along with limb and digit abnormalities (Bonilla-Claudio et al., 2012; Ozekin et al., 2020; Sacco et al., 2015; Suzuki et al., 2009).

In zebrafish, the long fin mutants (*lof, alf,* and *schl*), illustrate the vital significance of potassium channel conductance and electrophysiological signals in establishing accurate body-to-appendage proportions (Lanni et al., 2019; Perathoner et al., 2014; Stewart et al., 2021). While the mechanisms mentioned above highlight the involvement of potassium channels in regulating zebrafish fin development, our transcriptomic analysis revealed a progressive increase in the expression of *kcnk5b* over time following muscle contusion injury, indicating its role in natural healing. This discovery, along with the increased magnitude of heterotopic bone formation observed after pectoral fin injuries in the *kcnk5b* gain-of-function mutants, strongly point towards a potential role for Kcnk5b in regeneration following injury, influencing the extent of new bone formation. This not only opens up new avenues for investigating the interplay between potassium channels and the pathways governing regeneration but as these channels are often targets of small molecule drugs, its action also provides a basis for therapeutic targeting of MOT – one that is titratable.

We next investigated processes sufficient to promote heterotopic bone growth. Upon injury, regenerative inflammation promotes tissue repair by a timed and coordinated infiltration of diverse cell types and the secretion of growth factors, cytokines and lipids mediators (Caballero-Sanchez et al., 2022). One of the key aspects of this process is the release of interleukins, especially the Il-6 type cytokines which act locally and systemically to generate a variety of physiologic responses (Jawa et al., 2011). In multiple transcriptome datasets from regenerating tissues, including zebrafish, Il-11 has been demonstrated to be induced as a response to injury. We also noticed both paralogous genes encoding Il-11 (*il11a* and *il11b*) to be highly expressed 24 hours after the initial muscle contusion injury, followed by a decline in expression at subsequent time points. Consequently, we intended to investigate whether alterations in interleukin 11 expression within the inflammatory microenvironment could influence heterotopic bone formation response to injury. The adult viable homozygous loss-of-function zebrafish *il11ra* mutants were utilised for this purpose. Upon exposing these mutants to multiple injuries in the pectoral fin region, it was evident that the heterotopic bone formation response was significantly reduced compared to wild-type fish.

Studies have shown the upregulation of Il-11 following fractures and its role in promoting regeneration (Kidd et al., 2010). Additionally, transgenic mice overexpressing human IL-11, experience increased bone formation without an effect on bone resorption or osteoclastogenesis (Kidd et al., 2010), due to its enhancing the effect of BMP-2 and inhibiting bone marrow adipogenesis (Takeuchi et al., 2002). These observations indicate that Il-11 promotes osteoblastic activity at injury sites without influencing osteoclastic activity. This could clarify the decrease in bone volume observed in the injured fin of *il11ra^-/-^* mutants, potentially due to bone resorption by osteoclasts following the microfractures while the osteoblastic activity is impaired. Nevertheless, this discovery highlights a potential new path for intervention, through broadly targeting the inflammatory response soon after damage/surgery. Additionally, drugs targeting Il-11 can be utilized and could hold promise as a potential therapeutic strategy for mitigating heterotopic bone formation.

## Conclusion

Our investigation unveils for the first time in zebrafish, the occurrence of heterotopic bone formation in response to injuries, akin to the observed phenomenon in humans. The cumulative observations of heterotopic bone formation in the pectoral fin and caudal peduncle region, and consistent extensive proliferation and increase in size of the intermuscular bones following injury suggest that osteo-induction and proliferation of ectopic bone can occur in the zebrafish if appropriate injury methods are employed. Furthermore, the ability to induce consistent changes in bone structure after injury, and the subsequent imaging options available, it is without a doubt that zebrafish holds the potential to serve as a powerful and dependable experimental model for further research into heterotopic ossification. Employing this model, we observed a significantly increased magnitude of heterotopic bone formation in zebrafish with a gain-of-function mutation in the Kcnk5b channel. This finding suggests a central role for potassium channel signalling, particularly through Kcnk5b, in regulating the skeletogenic injury response. In contrast, zebrafish carrying a loss-of-function mutation of the Interleukin 11 receptor paralogue (Il11ra), known to be associated with impaired regeneration, exhibited a substantially reduced response. This contrast underscores the importance of interleukin signalling via Il11ra in the injury response mechanism leading to heterotopic bone formation. These findings not only advance our understanding of the molecular basis of heterotopic bone formation but also provide potential insights for therapeutic strategies for human patients grappling with this debilitating condition.

## Materials and Methods

### Zebrafish Husbandry

Zebrafish work was conducted at the fish facilities of Lee Kong Chian School of Medicine (LKCMedicine), Nanyang Technological University (NTU), Singapore and Boston Children’s Hospital (BCH), USA. All experimental procedures involving fish adhered to AAALAC standards and were granted approval by the Institutional Animal Care and Use Committees (IACUC) of NTU under protocol number A19028, and by BCH IACUC under protocol number 00001704. AB wild-type fish at the LKCMedicine fish facility were used to establish the injury model, perform RNA sequencing for transcriptomic analysis, and subsequent validation of the results. Furthermore, *Tg(sp7:egfp; ctsk:dsred)* reporting osteoblasts in green and osteoclasts in red were utilised for live imaging. *kcnk5b* mutants, *il11ra* mutants, and their wild-type sibling controls housed at the BCH fish facility were used for later experiments. All fish were maintained in facility water at 28°C, adhering to a 14-hour light and 10-hour dark cycle.

### Injury experiments

The contusion setup comprised a 45° wedge and a customized lancing device, with the depth setting typically adjusted to less than 1 mm, all positioned on a petri dish (Figure S5). The lancing device’s sharp tip was replaced with a custom epoxy resin spheroid to prevent piercing the skin, instead causing a blunt injury. Adult zebrafish of over 3 months post-fertilization were anesthetized with 0.013% Tricaine (buffered to pH 7.0). Following anaesthesia, the fish were placed laterally on the petri dish, ensuring that the target area rested on the impactor tip, and then subjected to 5-10 lancet strikes of similar force to induce a visible contusion. Early resolution of the contusion was addressed by gently stroking the injury site with a blunt-tip K-pin 48 hours after the first injury, reverting it to its post-1st injury state. This process was repeated, allowing the contusion phase to be sustained for up to a week. Regarding thoracic contusions, the lancing device setup was not employed for the initial injury; rather, stroking the region with the blunt-tip K-pin resulted in similar contusions as in the caudal peduncle region.

For pectoral fin injuries, following anaesthesia, fish were positioned on a petri dish in a lateral orientation with right side facing up. Subsequently, the pectoral fin was raised to expose the medial muscle bulk, and gentle stroking using a #5 Dumont forceps was carried out at the mid-section of this muscle bulk, guided by tactile feedback from the forceps’ tip as it grazed over the bone’s surface. This process resulted in visible contusions and microscopic damage to the pectoral fin rays. Both procedures were carried out under a Leica MZ7.5 dissecting stereomicroscope. No signs of distress were observed post-injuries in any of the fish.

### Alizarin red staining and imaging

Alizarin red staining of euthanised fish to visualize the skeleton was done based on the protocol described by TF Schilling (Schilling, 2002). Brightfield imaging was done using Axio Zoom. V16 (Zeiss) stereo zoom microscope. Measurements of bones, when necessary, were conducted utilizing Zen Blue 3.3 or Zen Lite 2.3 image analysis software by Carl Zeiss.

### RNA Sequencing

Tissues were harvested from the caudal peduncle site before and after injury from different sets of fish. Injuries were done in a similar manner as described above. Each condition i.e. No injury, 24 hours after 1 injury, 24 hours after 3 injuries, and 5 days after 3 injuries had three replicates. Each replicate was from 3 adult fish, approx. 0.5 cm muscle tissue from the caudal peduncle region proximal to the tail fin, taken after removal of the skin. Harvested sample was predominantly muscle tissue except for the tiny intermuscular bones that could be present. RNA was extracted as per previously published protocol (Peterson and Freeman, 2009). Initial assessment of its concentration and purity was conducted using a NanoDrop spectrophotometer. The RNA Integrity Number (RIN) was subsequently assessed with an Agilent 2100 Bioanalyzer, confirming that all samples met the necessary quality standards for sequencing. The mRNA library preparation and paired-end PE150 sequencing was carried out on Illumina’s Novaseq-6000 platform by NovogeneAIT Genomics Singapore Pte Ltd. Following quality control and pre-processing, the clean reads were mapped to the Zebrafish Genome Assembly GRCz11 (GCA_000002035.4) using *salmon* (v.1.9.0) (Patro et al., 2017), considering decoys and GC biases for zebrafish with an average of 83.8% (Range: 82.1-86.1%) fragment hits. All downstream data analyses were carried out in R/RStudio. Gene expression levels under different conditions were noted and the correlation of gene expression levels between the 12 samples was assessed using the Pearson’s correlation coefficient. While replicates had high similarity (R2 > 0.97) showing sample reproducibility, differential expression was noted between the 4 conditions (R2 range = 0.74 – 0.91). The *DESeq2* package (v.1.38.3) was used for differential analysis of count data, where lowly expressed counts (< 10) were filtered out from the dataset (Love et al., 2014). The *clusterProfiler* (v.4.6.0) package was used for Gene Ontology and KEGG pathway analysis (Wu et al., 2021). All figures were generated using *ggplot2* (Valero-Mora, 2010).

### qRT-PCR

For qRT-PCR validation, cDNA conversion was done using 1µg of DNase treated RNA sample, Oligo dT (Thermo Fisher Scientific), MultiV Reverse Transcriptase (RT) enzyme (New England BioLabs; NEB), 10x RT buffer (NEB) and 10mM dNTP mix (NEB). All qPCR reactions were carried out on an Applied Biosystems Step One plus real-time PCR system using standard thermal-cycling conditions optimised for KAPA SYBR® FAST qPCR Mix. The primer sequences used were as follows: *kcnk5b* forward (ATCACTCTCCTCGTCTGCAACG) and reverse (GAGTCCCATGCACAACGTGCAG); *il11a* forward (GGACAAATATGAAATTGCTGGGTG) and reverse (AGCGTCAGAAGGAGTTTGGT); *il11b* forward (TGAACGCAAATGAGTTGACTG) and reverse (CCCAATTCGTCACTATTCCGT); *rpl13a* forward (AGACGCACAATCTTGAGAGCAG) and reverse (TCTGGAGGACTGTAAGAGGTATGC); *dlx3b* forward (TGAGTGGACCGACATATGACA) and reverse (TTAATACACGGCCCCCACG); *col10a1a* forward (CCTGGAGCCAAAGGAGAGTT) and reverse (TATCGGCAGCAAAGACACCA); and *mmp9* forward (CAGTGGAAATGATGTGCTTGG) and reverse (GAAGTAGAAGAATCCCTTGTAGAG).

Data was exported as excel files and analysed using the 2^-ΔΔCT^ method (Livak and Schmittgen, 2001).

### Micro-CT scan

Micro-CT scans were performed on injured and uninjured pectoral fins of *kcnk5b and il11ra* mutants, as well as their wild-type controls. After routine alizarin red staining and two-dimensional imaging, the fins were stained with 1% silver nitrate in multi-well cell culture dishes placed on a standard gel lightbox for a duration ranging from 45 minutes to an hour. The actual duration was determined based on the degree of staining observed in fin rays and the background intensity. After adequate staining, the samples were washed with distilled water three times and then fixed onto agar blocks and stored at 4°C until scanning. Sanning was done using Skyscan 1173 micro-CT scanner (Bruker). Following the scan, volume-rendered images of the contrast-stained samples were created using the Amira software package, version 6.0 (Thermo Fisher) and saved as nifti files. Subsequent, volumetric assessments were done using 3D slicer (Kikinis et al., 2014).

### Statistical analysis

All statistical analyses were performed using R and GraphPad Prism 9 (GraphPad Software Inc, San Diego, CA), as deemed appropriate. Before comparing continuous variables, the assumption of normal distribution was assessed using the Shapiro-Wilk test. In cases where the data exhibited a normal distribution, a student’s t-test was employed for comparison. Conversely, when the data did not follow a normal distribution, the Mann-Whitney U test was utilized to ascertain statistical significance. For evaluating associations between categorical variables, Fisher’s exact test was employed. A p-value of less than 0.05 was considered statistically significant, unless otherwise stated.

## Code and data availability statement

The FASTQ files of raw RNAseq data are openly available in NTU research data repository DR-NTU (Data) at https://doi.org/10.21979/N9/HEJN6X. The *shell* and *R* scripts to reproduce the RNASeq analysis can be found at https://github.com/cenk-celik/kaliya_perumal_d_rerio.

## Acknowledgements

We sincerely thank Dr Sven Reischauer for generously providing the *il11ra*^-/-^ mutant zebrafish. We also thank the McMenamin lab for assistance with performing the micro-CT scans. We are indebted to Azmi Bin Ja’afar, Joshua Gan Lan Teng and Neo Sui Hoon for maintenance of our zebrafish lines. This research was funded by the Toh Kian Chui foundation (award to PWI). AKKP was supported by a Nanyang Technological University Research Scholarship.

