## Supplementary material for "Genetic regulation of injury induced heterotopic ossification in adult zebrafish": Figure S1-S5

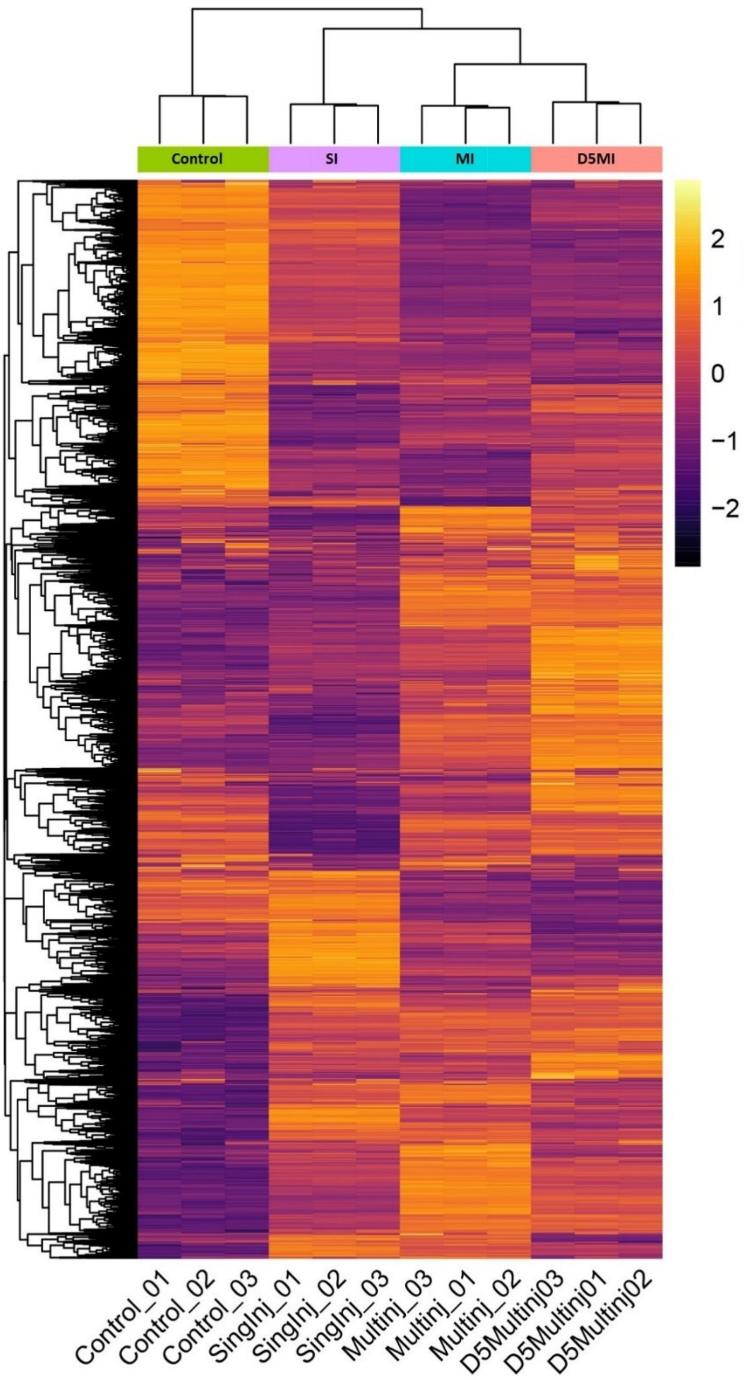


**Figure S1.** Heat map of the top 10,000 differentially expressed genes (DEGs), revealing a temporal relationship with expression profiles of the D5MI group more closely related to the MI group when compared to the SI group and Control group, which were the most distantly related.

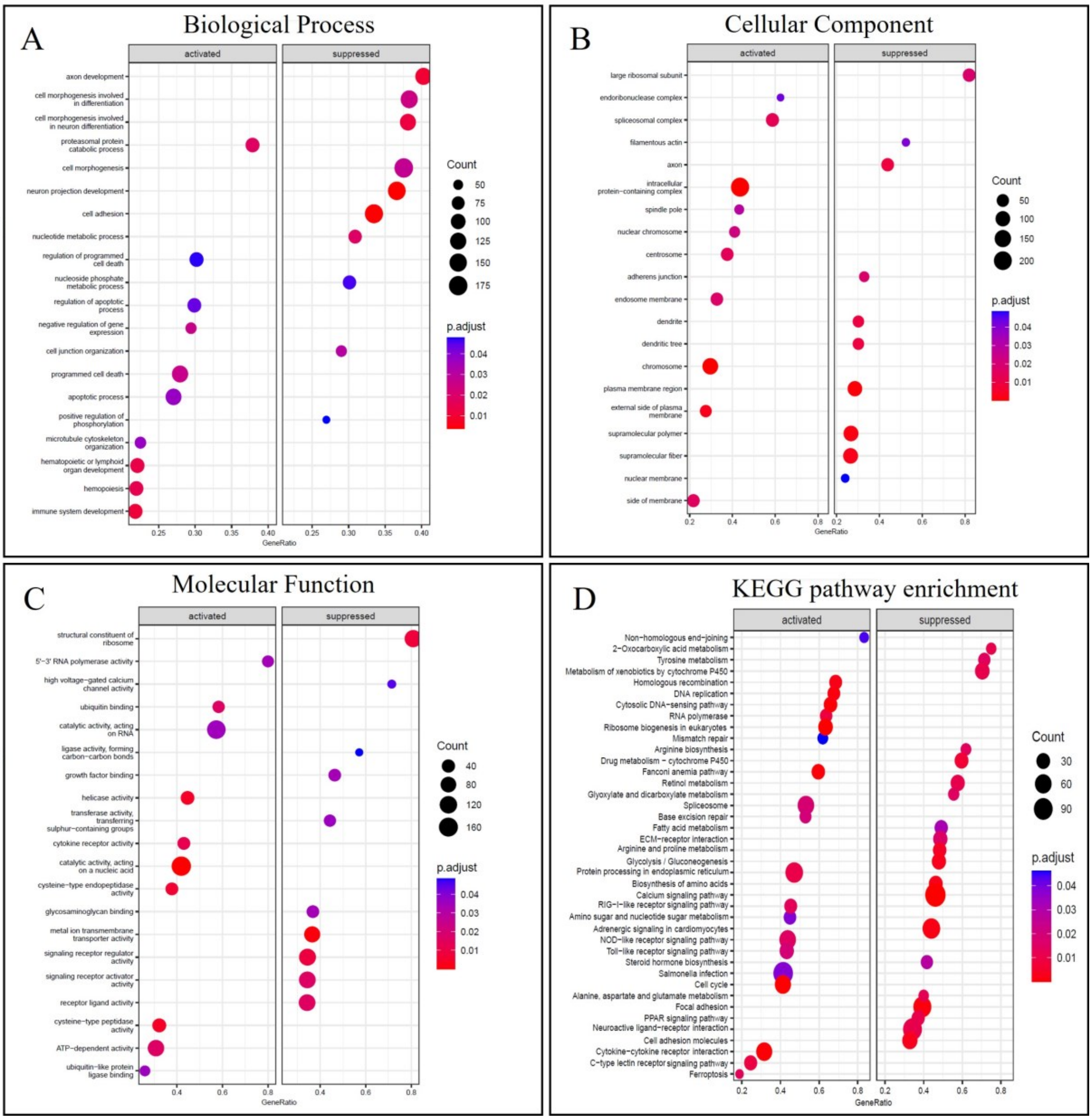


**Figure S2.** Functional annotations based on GO terminology and KEGG pathway enrichment for comparison 1 (Single injury vs. Control). **A)** Biological process. **B)** Cellular component. **C)** Molecular function and **D)** KEGG Pathway Enrichment.

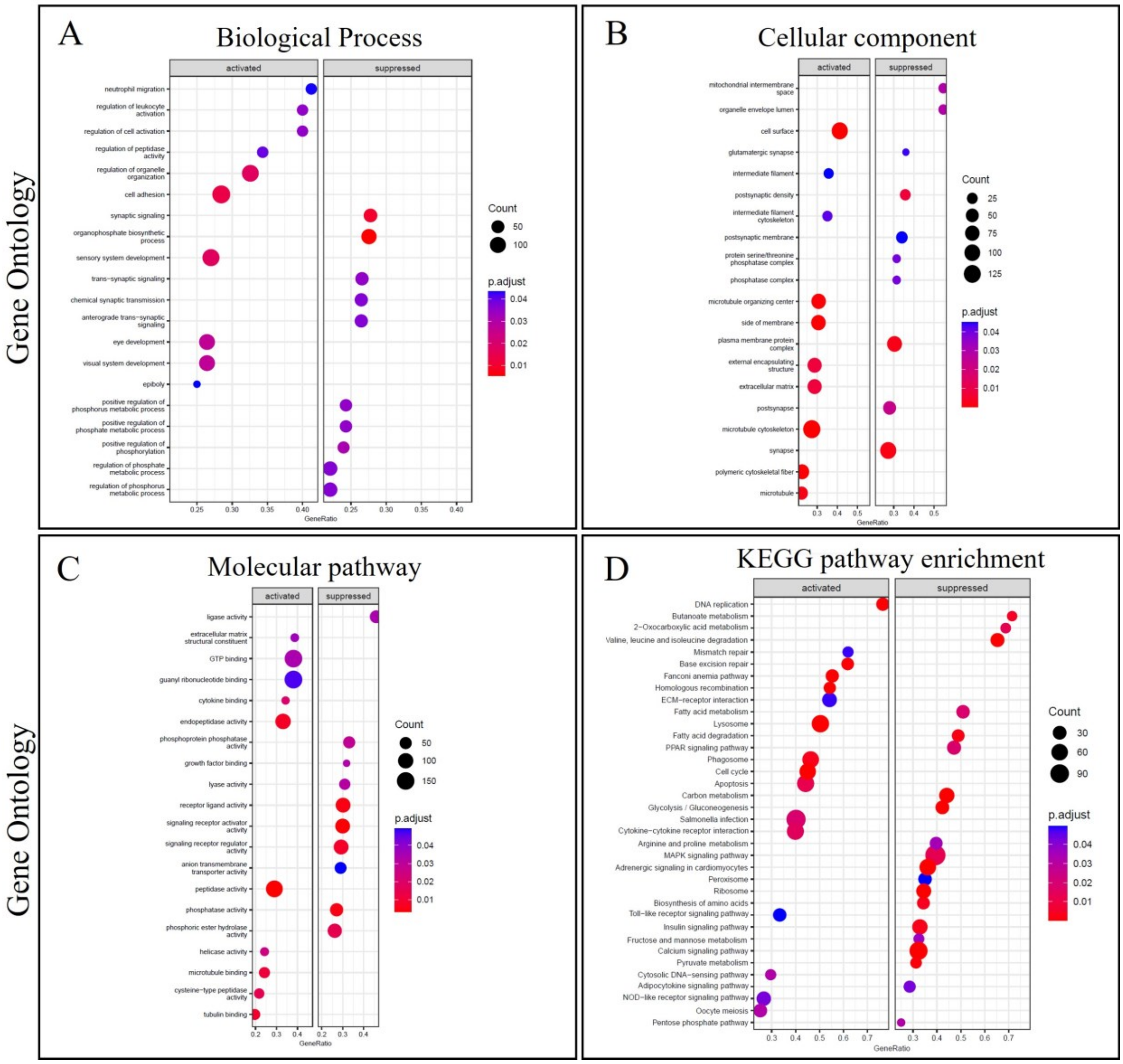


**Figure S3.** Functional annotations based on GO terminology and KEGG pathway enrichment for comparison 2 (Multiple injuries vs. Control). A) Biological process. B) Cellular component. C) Molecular function and D) KEGG Pathway Enrichment.

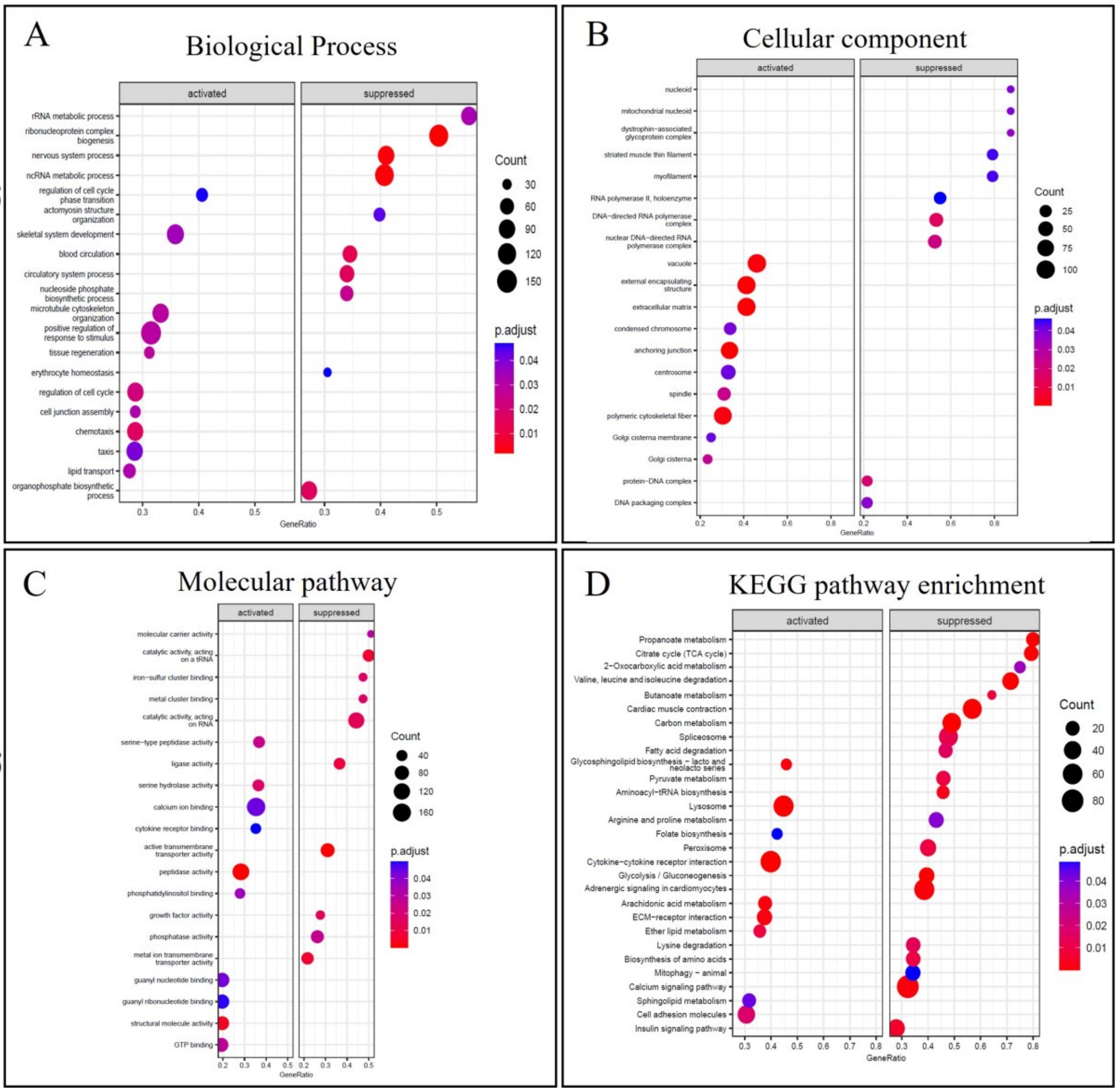


**Figure S4.** Functional annotations based on GO terminology and KEGG pathway enrichment for comparison 3 (5 Days after multiple injuries vs. Control) A) Biological process. B) Cellular component. C) Molecular function and D) KEGG Pathway Enrichment.

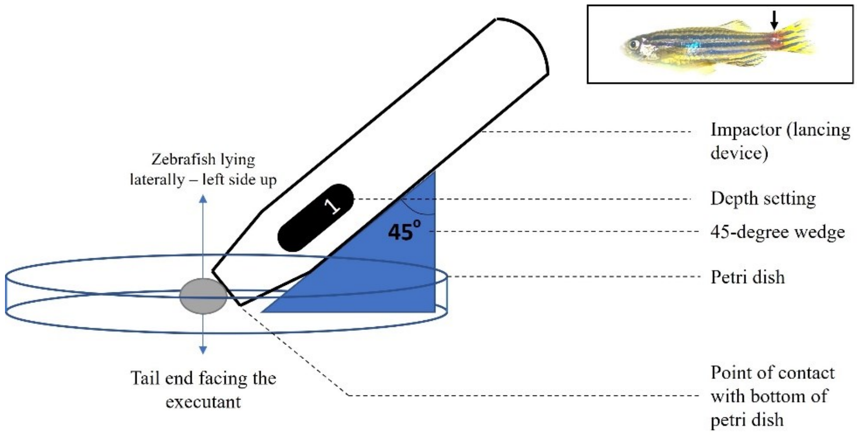


**Figure S5.** Set up for creating caudal peduncle contusion. Boxed area showing an adult zebrafish post contusion (arrow).

**Table S1:** Statistically significant differentially expressed genes for comparison 1 (Single injury vs. Control)

**Table S2:** Statistically significant differentially expressed genes for comparison 2 (Multiple injuries vs. Control).

**Table S3:** Statistically significant differentially expressed genes for comparison 3 (5 Days after multiple injuries vs. Control)

**Movie 1:** 3D reconstructed CT scan of an injured pectoral fin in $kcnk5b^{pfau+/-}$ mutant zebrafish, revealing extensive heterotopic ossification on the medial aspect.

**Movie 2:** 3D reconstructed CT scan of an injured pectoral fin in $il11ra^{-/-}$ mutant zebrafish, showing no signs of heterotopic bone.
